# Opportunities for prioritizing and expanding conservation enterprise in India using carnivores as flagships

**DOI:** 10.1101/2020.01.03.894311

**Authors:** Arjun Srivathsa, Iravatee Majgaonkar, Sushma Sharma, Priya Singh, Girish Arjun Punjabi, Malaika Mathew Chawla, Aditya Banerjee

## Abstract

Conservation interventions in developing countries are frequently thwarted by socio-economic agendas, severely limiting the scope and rigor of biodiversity and habitat conservation. Very few ecological assessments incorporate human interests in conservation prioritization, creating asynchrony between planning and implementation. For conservation actions to be logistically feasible, multiple criteria including ecological, social, economic and administrative aspects must be considered. Understanding how these different dimensions interact spatially is also important for gauging the potential for conservation success. Here, we use a guild of select mammalian carnivores (wild canids and hyenas) in India to (i) generate distribution maps at the spatial scale of administrative sub-districts, that is relevant to management, (ii) examine ecological, social and biogeographic factors associated with their distribution, quantify key threats, and identify areas important for their conservation, (iii) use prioritization tools for balancing habitat conservation, human needs and economic growth, and (iv) evaluate the spatial congruence between areas with high conservation potential, and areas currently in focus for protection efforts, conservation investments, and infrastructure development. We find that the current Protected Area system does not adequately cover or represent diverse habitats, that there is immense potential for States to increase financial investments towards alternative conservation strategies, and, most infrastructure projects may be jeopardizing important carnivore habitats. Our framework allowed for identifying locations where conservation investments would lead to the highest dividends for flagship carnivores and associated species across habitats. We make a case for re-evaluating how large-scale prioritization assessments are made, and for broadening the purview of conservation policies in India and other developing countries.

## Introduction

Conservation is a politically challenging endeavor, particularly in developing economies where governments have to frequently negotiate a balance between limited conservation funding and pressing economic agendas (Waldron et al. 2013; Lindsey et al. 2016). In this context, debates on what to conserve and how, the use of relevant biodiversity surrogates, and trade-offs in conservation planning are integral parts of the conservation discourse (Westgate et al. 2014; Lentini & Wintle 2015). The advent of decision-support tools have enabled scientists to quantitatively determine priority hotspots, prescribe ideal reserve designs, and better plan land-use management for conservation (Moilanen et al. 2005; Ball et al. 2009; Gregory et al. 2012). However, these aspects are rarely in tandem with on-ground implementation due to socio-political, cultural and economic hurdles (Game et al. 2013), seldom meeting requirements of representativeness and complementarity in species and habitat conservation (sensu Margules and Pressey 2000). As a consequence, allocation of research efforts, human resources and conservation funding in many countries do not agree with global priorities and goals for conservation (Halpern 2006; Jenkins et al. 2013; Brum et al. 2017). Given this background, conservation prioritization assessments, more often than not, become limited to an academic exercise.

Integrating federal research and decision-support tools with conservation interventions have been successful in select countries and landscapes, mostly in the global north (Wintle et al. 2019). The global south, on the other hand, exemplifies certain asynchronies in priorities, where balancing human well-being, economic development, and safeguarding nature at the same time, present formidable challenges (Sandker et al. 2012; Barlow et al. 2018). Many developing countries in the tropics have witnessed rapid loss of species and habitats, accounting for some of the highest extinction rates in the world (Pimm et al. 2014; Ceballos et al. 2017). But these tropical countries are also investing heavily on infrastructure development, relying on capitalist approaches to economic growth, often at a cost to natural habitats and threatened species (Barlow et al. 2018). The expanding and growing economy in these countries, together with a globalizing human society aspiring for better standards of living, continue to sustain this demand. Such scenarios call for multi-criteria conservation planning assessments that stretch beyond taxonomic, phylogenetic or ecological considerations (Brum et al. 2017; Sibarani et al. 2019) to additionally acknowledge, address and incorporate socio-economic, political and administrative factors.

Globally, Protected Area networks are dominated by forested biomes, with 2–6 % of tropical, subtropical and temperate savanna grasslands and shrublands afforded strict protection (Jenkins & Joppa 2009). Most published literature and policy perspectives deal with prioritizing global species-rich hotspots in tropical forests, or Protected Areas and woodland systems in relatively wealthy, temperate countries (Martin et al. 2012). Furthermore, management approaches that decouple human-use areas and wilderness areas in pursuit of “intactness” (Mittermeier et al. 2003), tend to bias protection efforts towards landscapes with very low human densities or access (Joppa and Pfaff 2009; Venter et al. 2014). Such approaches undervalue many ecosystems and species that do not fit the parochial view of nature and wilderness. For example, savanna grassland systems, semi-arid scrublands, ravines, deserts and pasturelands may not have very low human densities, but they support unique assemblages of species adapted to survive in such ecosystems (e.g., Brito et al. 2013). Even cultivated agricultural lands support many large mammalian species that persist in transformed food webs and systems (Magioli et al. 2019). Such multi-use areas rarely figure in global priority locations for conservation (Jenkins et al. 2013), whilst being lost at an alarming rate or rapidly transitioning into unsustainable use (Dobrovolski et al. 2013). Basic ecological information on key indicator species that represent the health of these habitats is severely inadequate or inaccurate (Hurlbert & Jetz 2007; Ghosh-Harihar et al. 2019). This creates an imperative need for identifying and using appropriate flagship species (Verissimo et al. 2013; Shen et al. 2019) or species assemblages as surrogates to conserve such diverse yet neglected ecosystems.

Terrestrial mammals that belong to the order Carnivora receive among the highest investments in terms of conservation funds, human resources and public support (Treves & Karanth 2003; Smith et al. 2012). That some of the most admired species worldover are large wild felids further testifies their charismatic appeal (Macdonald et al. 2015). Mammalian carnivores represent a spectrum of habitat associations (Crooks et al. 2011; Bateman & Fleming 2012), with some showing extremely high specificity (e.g., Ethiopian wolf *Canis simensis* in the highlands of Ethiopia) and others representing a range of eclectic land cover types (e.g., golden jackal *Canis aureus*). Certain predators like the leopard *Panthera pardus*, coyote *Canis latrans* and puma *Puma concolor* even thrive in radically human-modified habitats, sometimes sharing space with high human densities (Gehrt & Riley 2010; Vynne et al. 2011; Athreya et al. 2015). Recent studies have also highlighted the variety of critical ecosystem services and benefits carnivores provide, all of which contribute towards human well-being through direct or indirect mechanisms (O’Bryan et al. 2018). Yet, carnivores are highly threatened across the world, with populations facing massive range contractions and risks of local extinctions, primarily due to anthropogenic factors (Ripple et al. 2014). Taken together, their natural capital, threatened status, and the ecological and trophic niches they occupy, makes carnivores potential flagships in their respective habitats for targeting conservation efforts (Macdonald & Loveridge 2010).

Here, we use India as a case study and focus on nine species/sub-species (henceforth ‘species’) of carnivores– dhole *Cuon alpinus*, golden jackal, Indian wolf *Canis lupus pallipes*, Tibetan wolf *Canis lupus chanco*, Indian fox *Vulpes bengalensis*, red fox *Vulpes vulpes*, desert fox *Vulpes vulpes pusilla*, Tibetan fox *Vulpes ferrilata* and striped hyena *Hyaena hyaena* (Fig. 1). We:

i. generate countrywide distribution maps of the carnivores, at a spatial scale and resolution that is relevant to administration
ii. examine ecological, social and biogeographic factors associated with their distribution, quantify key threats, and identify landscapes and habitats critical for their conservation
iii. use prioritization tools to make a case for conserving a consortium of habitats with wild canids and hyenas as flagships, while balancing human needs and economic development
iv. evaluate the spatial congruence between areas with high conservation potential, and areas currently in focus for protection efforts, conservation investment, and infrastructure development.

**Figure 1.**
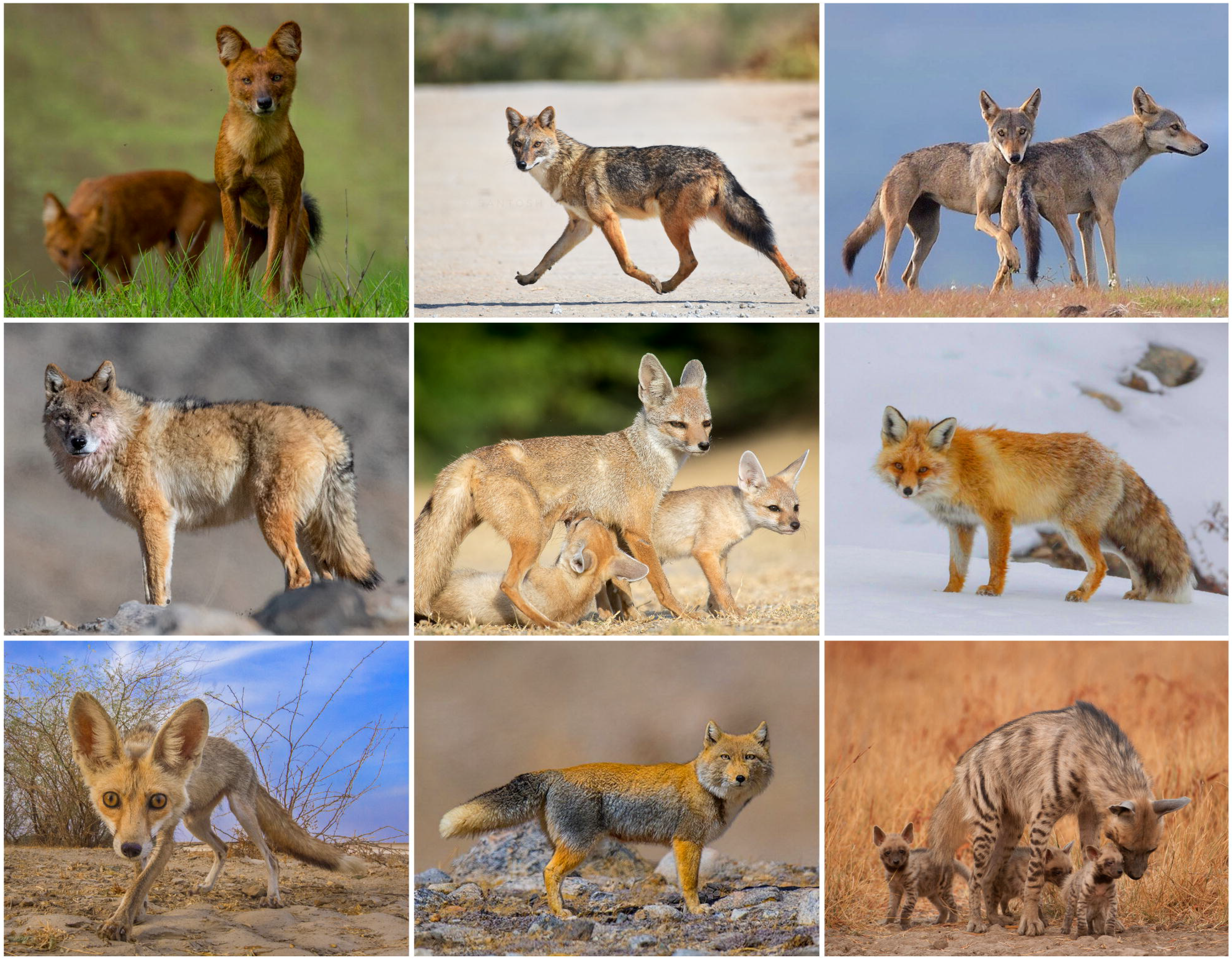
The nine focal carnivore species/sub-species in this study. From left to right, top row to bottom row (photo credits in parentheses): dhole (Sushvin Gowda), golden jackal (Santosh Doddagoudar), Indian wolf (Anup Deodhar), Tibetan wolf (Aditya Chavan), Indian fox (Anuroop Krishnan), red fox (Ismail Shariff), desert fox (Manish Vaidya), Tibetan fox (Aditya Chavan) and striped hyena (Kalyan Varma).

In doing so, we provide a roadmap for implementing a decentralised approach to long-term monitoring of carnivores and the habitats they represent, through periodic assessments of their distribution patterns, threats, human interests and administrative management potential.

## Materials and Methods

### Carnivore presence records

We collected carnivore distribution data in three phases. In phase 1, we used citizen-science data from countrywide web-based surveys for three months in 2018 (see SI Appendix, section 1 for detailed survey protocol). In addition to our own survey, we also included carefully vetted records from other citizen-science portals (India Biodiversity Portal: http://www.indiabiodiversity.org; iNaturalist: https://www.inaturalist.org). In phase 2, we extracted data from wildlife, nature and photography pages on social media, online wildlife photo-repositories and reliable blog articles in 2018–19 (SI Appendix, section 1). In phase 3, we extracted information from published studies, unpublished theses, forest department reports and openly accessible project reports submitted to funding agencies in 2019. We thoroughly verified and validated each record, ensuring correct species identification, geographic location, and time (month and year) of record. We considered only reliable records of species presence, corresponding to the period from January 2015 to December 2018 (SI Appendix, section 1).

### Distribution models

We treated administrative sub-districts (‘*tehsil*’ or ‘*taluk*’) in the country as independent spatial units or ‘sites’ (mainland India has 2342 sub-districts). We chose sub-districts as our units because this spatial scale provided a balance between being ecologically meaningful and also relevant for implementing management actions. For each species, we first defined a plausible distribution range based on all available information from field guides, State forest department checklists, published literature and our own field-based knowledge. Every presence record was then assigned to an administrative sub-district within the species’ plausible range. We used an occupancy modelling framework that accounted for partial detectability to map distribution patterns (MacKenzie et al. 2018). Since our data pertained to presence-only information, we created detection histories using the following method– for each species, a given site within its plausible range was labelled ‘D’ (detected) if it had at least one detection in the four-year period; months with detections were assigned ‘1’ and months with no detections were assigned ‘0’. Sites were labelled ‘ND’ (not detected) when the focal species was not detected across four years, but there was at least one detection of any of the other eight species; all months in these sites were assigned ‘0’. Sites that did not have detections of any of the nine species during the four-year period were labelled ‘NS’ (not surveyed; see Powney et al. 2019). For analyses, we collapsed data from 48 months into four 12-month blocks, resulting in one temporal replicate per year. At the sub-district scale (mean = 1400 sq. km; range = 3 – 51,000 sq. km), we assumed that the distribution status of the focal species remained stable during the four-year period. For each species, we built a set of candidate models with singular and additive effects of a set of explanatory variables, based on specific *a priori* predictions (SI Appendix, section 2). We fit occupancy models to detection/non-detection data using package ‘unmarked’ in program R v3.4.1 (R Core Team 2018).

### Explanatory variables for modeling distributions

We used a combination of remotely sensed data, government-generated figures, and estimates from published studies to compile information on explanatory variables. These variables were chosen based on their expected influence on one or more species. We used 12 land-use land cover categories, synthesized and combined from a total of 152 categories classified by Roy et al. (2015). Climate and topography data were extracted from remotely sensed satellite imagery. Data on Protected Areas, linear infrastructure (railways/roadways) and human population densities were obtained from web-based open data sources. Population data on livestock (sheep, goat and cattle) and free-ranging dogs were sourced from government livestock census, and data on large wild prey (for dhole and Indian wolf) were based on published literature. All the variables were re-processed at the sub-district scale for analyses. Details of these variables, including category, description, and data source and year are in Table S1.

### Threat assessment

Given the paucity of quantitative information on human-induced threats to wild canids and hyenas in India, we created a conservation score for each species by combining their protection status, area of occupancy in India, and threat information from surveys of field experts. For protection status, we included the IUCN Red List status (scores: 1 = CR; 2 = EN; 3 = VU; 4 = NT; 5 = LC), CITES Appendix category (5 = Appendix 1; 3 = Appendix 2; 1 = Appendix 3), and India’s Wildlife Protection Act Schedule (5 = Schedule l; 3 = Schedule 2; 1 = Schedule 3). Scores for area of occupancy were based on the predicted proportion of habitat that a species currently occupies within its plausible range in India (scores: 1 to 10; 1 = species occupies 0–10% of plausible range; 10 = species occupies 90–100% of plausible range). Through surveys of field experts, we elicited scores for perceived population trend (1 = decreasing; 3 = stable; 5 = increasing). We also obtained scores for ‘level of threat’ attributed to habitat loss, prey decline, direct persecution, road-related mortality, illegal trade and negative interactions with free-ranging dogs (1 = fatal; 2 = high; 3 = medium; 4 = low; 5 = not a threat). The total scores for each category were averages from scores of individual experts. The final conservation score was obtained by summing across all categories, weighting area of occupancy at 0.5, protection status at 0.3 and expert responses on threats at 0.2. The unequal weighting is because area of occupancy is a quantitatively estimated metric, protection status is derived from global datasets but without the same analytical rigor, and threat information is based on expert opinions (which could be anecdotal, or limited to insights from local/regional experience). We re-scaled the sum out of 100; a lower conservation score implied a higher threatened status (within India).

### Spatial prioritization

We used software Zonation 4.0 (Moilanen et al. 2014) to identify regions of high and low conservation values with respect to the focal carnivore species and their habitats in India. Zonation’s algorithm ranks cells (raster pixels) within the area of interest through an iterative cell-removal process. This is implemented by first assuming that all cells are to be retained, and then, removing cells that cause the smallest marginal loss in the overall conservation value (Moilanen et al. 2014). The process is repeated until no more cells are left; cells with the least value are removed first and those with the highest values are retained towards the end. The relative weights assigned to the input features and the cell-specific value of each feature determines the hierarchy of cell removal. The output raster file thus includes pixels ranked by relative conservation priority values. We chose the ‘Core Area Zonation’ cell removal rule because it prioritizes units with the most important and rare features while minimizing loss in the overall conservation value. We also modified the algorithm such that the final conservation values are assigned to clusters of pixels grouped by sub-district boundaries. As primary input features, we used (i) carnivore diversity indices calculated from estimated occupancy probabilities, and (ii) extent of important habitats in each sub-district (Table S2). We chose diversity indices rather than individual species distributions after preliminary exploratory analyses showed better spatial representation of important areas and greater area of retention with the former. Diversity indices used as input features are shown in Fig. S1. In addition to these, we also included human poverty index and projected human population for 2020, both of which qualified as “costs” in our assessment. Our rationale was that sub-districts with larger human populations and higher poverty require focus on economic growth and infrastructure development and should therefore be of lower priority for conservation (see Table S2 for details). Following exploratory runs with different combinations of settings for Boundary Length Penalty (BLP) and Warp Factor (WP), we set BLP at 0 and WP at 100 as a trade-off between computation time and reliability of spatial maps.

Finally, we compared locations of high conservation priority and (i) their spatial congruence with current Protected Areas, (ii) State-wise conservation likelihood based on current budgetary spending in forest and wildlife sectors (because conservation investments are determined, sanctioned and disbursed by the State government), (iii) their overlap with locations of current/planned infrastructure projects such as solar farms, large- and small-hydropower projects, large mines, and irrigation projects– all of which are likely to affect the focal species and their habitats through habitat loss/fragmentation, either directly or through ancillary impacts.

## Results

We collated a total of 4437 presence records of the target species across three phases (Table 1). Model-averaged occupancy estimates ranged from 0.21 (SE 0.02) for desert fox to 0.75 (SE 0.002) for golden jackal (Table 1; Fig. 2). We could not formally analyze or generate estimates for Tibetan fox because the data were too sparse. Tibetan wolf occupied the least overall area (152,180 sq. km) and golden jackal was the most widely distributed species (2,259,361 sq. km). Combining protection status, distribution extent, population status and anthropogenic threats (the latter two elicited from surveys of field experts in India; n = 45), golden jackal and red fox scored the highest, while Indian fox and striped hyena scored the least (Table 1).

**Table 1.** Summary of data records collated from three survey phases, estimated occupancy (standard errors in parentheses), extent of occurrence in India and conservation score for the focal carnivore species

| Species | Phase 1 records | Phase 2 records | Phase 3 records | Estimated occupancy (SE) | Area occupied (sq. km) | Conservation score (scaled to 100) |
| --- | --- | --- | --- | --- | --- | --- |
| dhole | 290 | 417 | 82 | 0.46 (0.01) | 249,606 | 59 |
| golden jackal | 1054 | 377 | 86 | 0.75 (0.002) | 2,259,361 | 69 |
| Indian wolf | 331 | 177 | 8 | 0.33 (0.003) | 293,947 | 60 |
| Tibetan wolf | 92 | 60 | 1 | 0.36 (0.01) | 152,180 | 66 |
| Indian fox | 418 | 213 | 1 | 0.34 (0.001) | 890,831 | 49 |
| red fox | 58 | 54 | 95 | 0.50 (0.02) | 274,292 | 69 |
| desert fox | 28 | 115 | 0 | 0.21 (0.02) | 174,580 | 55 |
| Tibetan fox | 3 | 10 | 0 | — | — | — |
| striped hyena | 226 | 185 | 56 | 0.41 (0.001) | 957,478 | 49 |

**Figure 2.**
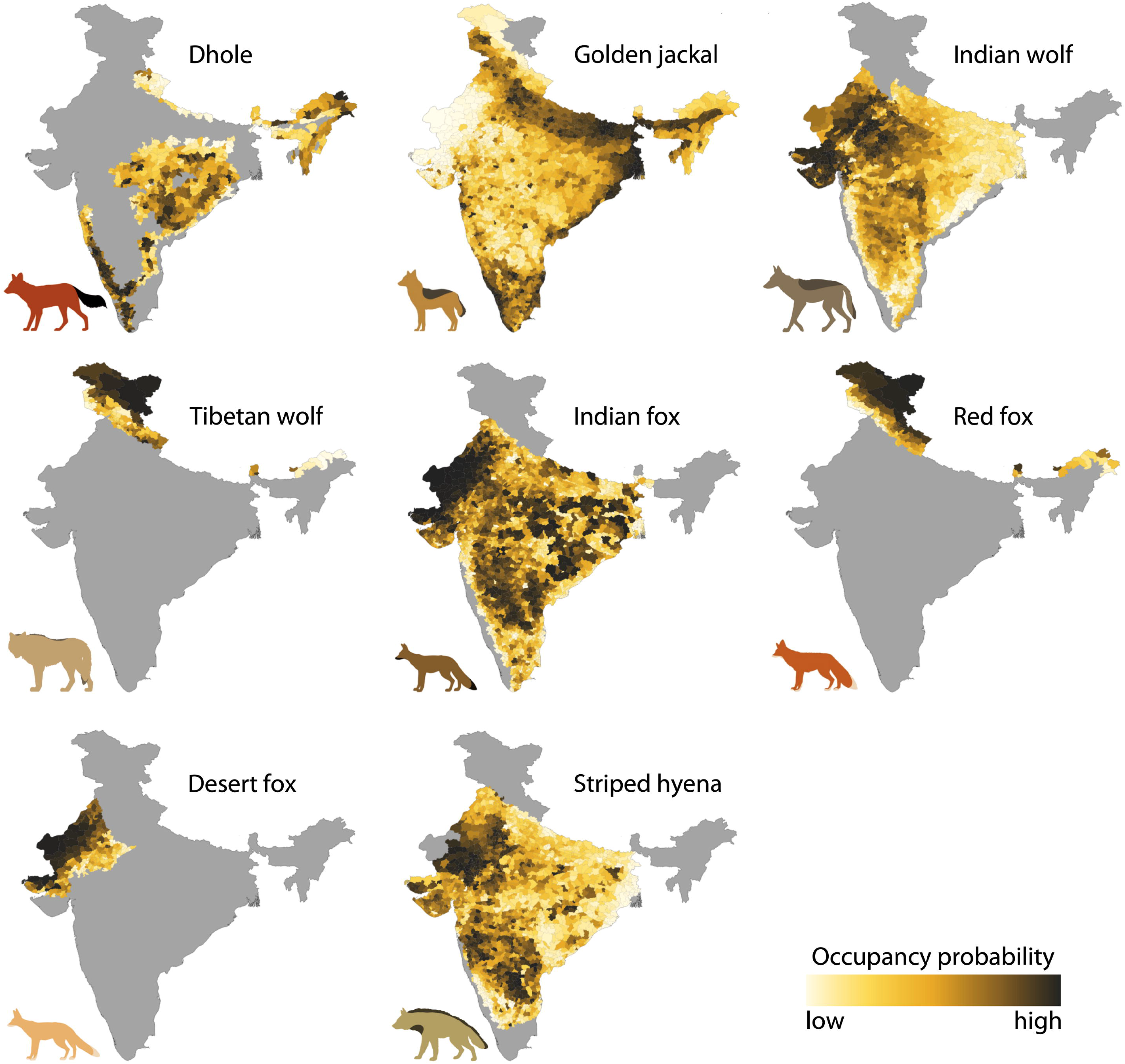
Estimated sub-district level occupancy probabilities for the focal carnivore species across India. Range of estimates: dhole (0.03–0.96), golden jackal (0.51–0.99), Indian wolf (0.02–0.55), Tibetan wolf (0.20–0.65), Indian fox (0.11–0.93), red fox (0.04–0.99), desert fox (0.02–0.99) and striped hyena (0.26–0.66). Model averaged estimates and associated standard errors are in Table 1.

Carnivore–habitat associations generally agreed with our predictions. Forest cover and large wild prey were important for dhole and Indian wolf (Table 2). Extent of grasslands, scrublands, ravines and open (barren) habitats were positively associated with all fox species and striped hyena. Production agroforests were preferred by dhole and jackal, and striped hyenas preferred rocky outcrops. Considered together, grasslands, scrublands, open/ravine habitats and forests were key habitats for wild canids and hyenas. Dhole occupancy was negatively associated with cattle and human densities, and positively with extent of Protected Areas– underscoring their importance for dhole source populations. The magnitudes and directions of effects with respect to human settlements, linear infrastructure and free-ranging dog densities varied across species (Table 2). Interestingly, high probabilities of golden jackal occurrence were clustered around large settlements with high densities of humans and free-ranging dogs. This may be attributed to high availability of provisioned food resources, and their ability to adapt to human-modified areas. While these results were based on singular covariate effects, the final occupancy estimates for all species were derived from averaging across models with comparable statistical support (based on AICc scores and weights; SI Appendix section 2, Table S3). We generated both, distribution maps at the sub-district level (Fig. 2), and maps depicting occurrence probabilities within plausible habitats (Fig. S2).

**Table 2.**
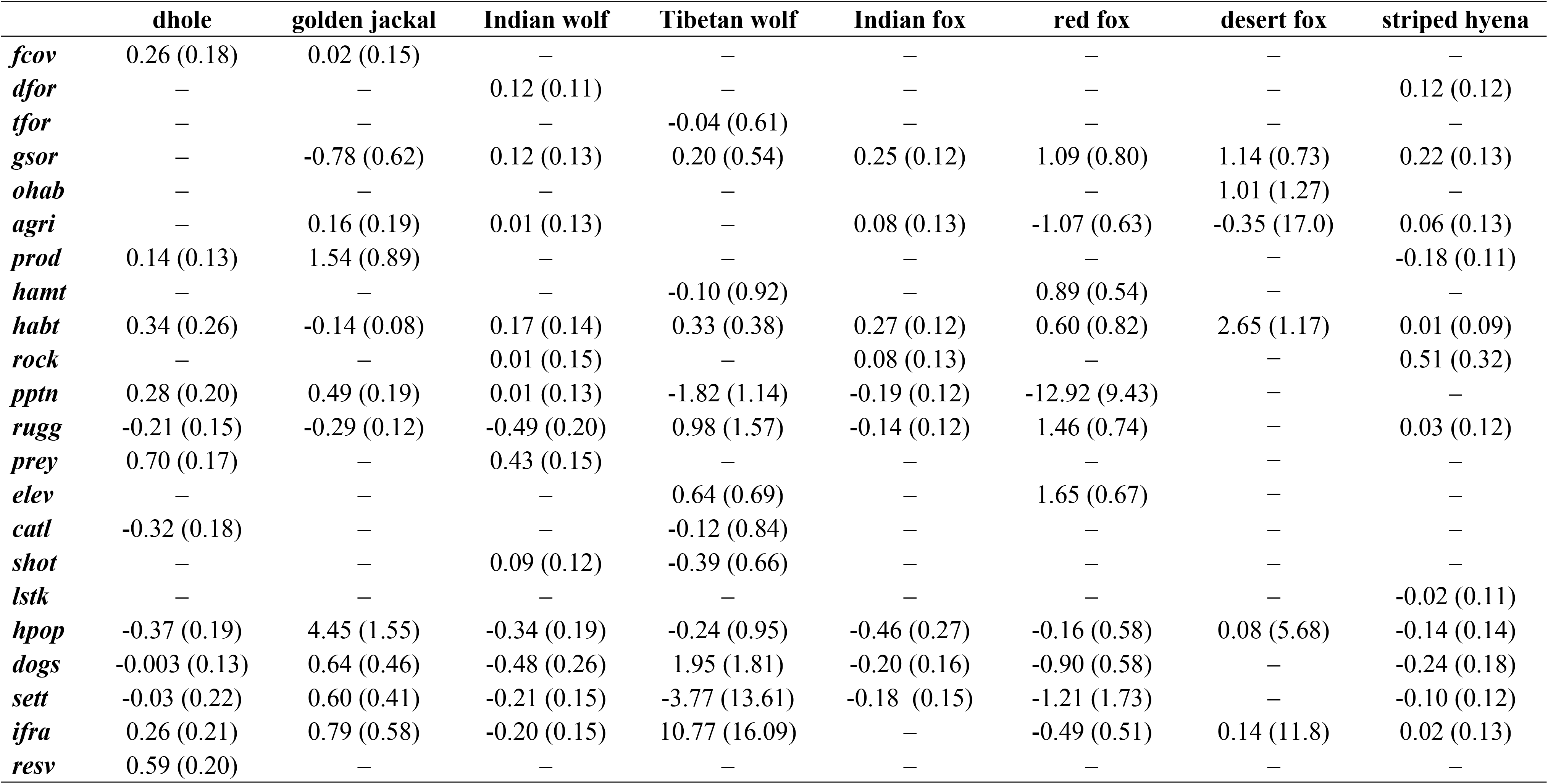
Estimated slope coefficients from singular covariate models for each of the carnivore species (standard errors in parentheses). Abbreviations: fcov– combined forest cover; dfor– tropical dry forests; tfor– temperate forests; gsor– grasslands, scrublands, open (barren) habitats and ravines; ohab– open (barren) habitats; agri– agricultural areas; prod– production agroforests; hamt– high altitude mountains; habt– combined area of plausible habitats (species-specific); rock– rocky outcrops and escarpments; pptn– annual precipitation; rugg– terrain ruggedness; prey– wild prey index; elev– elevation; catl– cattle population; shot– sheep and goat population; lstk– livestock (cattle, sheep and goat) population; hpop– human population; dogs– free-ranging dog population; sett– human settlements; ifra– density of roads and railways; resv– extent of Protected Areas

Based on results from the prioritization analysis, we chose sub-districts that accounted for the top 30% priority scores (n = 703) as the most important areas for conserving wild canids, hyenas, and their habitats (Fig. 3). The high priority areas– which we term ‘Canid Conservation Units’ (CCUs)– cover 1,475,558 sq. km, of which 545,070 sq. km (37%) constitute focal habitats (i. e., habitats deemed important for the focal species; Fig. 3). We further classified these areas into tier 1, tier 2 and tier 3, based on level of priority (top 10, 20, or 30% scores). Protected Areas currently cover around 26% of key habitats within CCUs (Fig. 3). Summed conservation ranks of CCUs showed that 12 States fared relatively better than India’s 17 other States (Fig. 4). Of these, Madhya Pradesh, Karnataka, Odisha and Chhattisgarh have the highest conservation potential and likelihood of success, when the States’ budgetary allocation for forests and wildlife sectors is taken into account (indicative of the States’ propensity to prioritize conservation of wildlife). Rajasthan, Gujarat, Andhra Pradesh, Maharashtra, West Bengal, Bihar, Tamil Nadu and Uttar Pradesh have high conservation scores, but will need to invest higher monetary resources to be able to work towards conserving CCUs in their States (Fig. 4).

**Figure 3.**
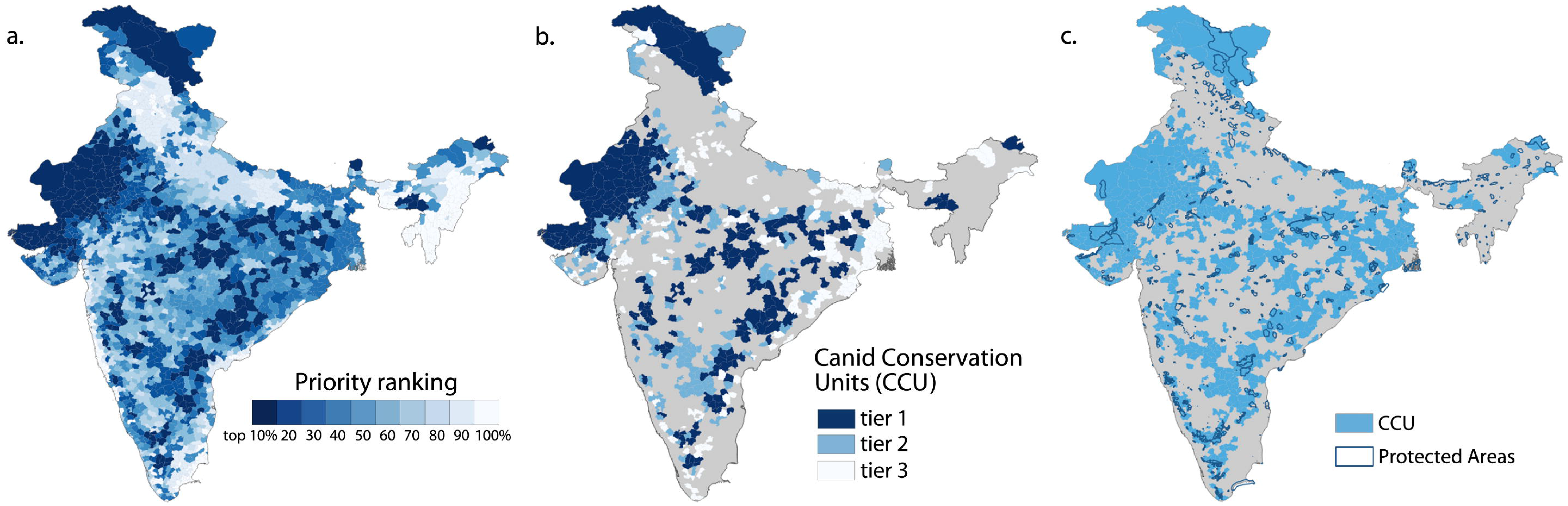
Spatial maps of results from prioritization analysis. (a) Sub-district level conservation priority scores for India. (b) Sub-districts with top 30% priority scores, classified as tiers 1– top 10% scores, 2– top 20%, and 3– top 30%, collectively termed Canid Conservation Units ‘CCUs’. (c) spatial overlap between Protected Areas of India and CCUs identified in this study.

**Figure 4.**
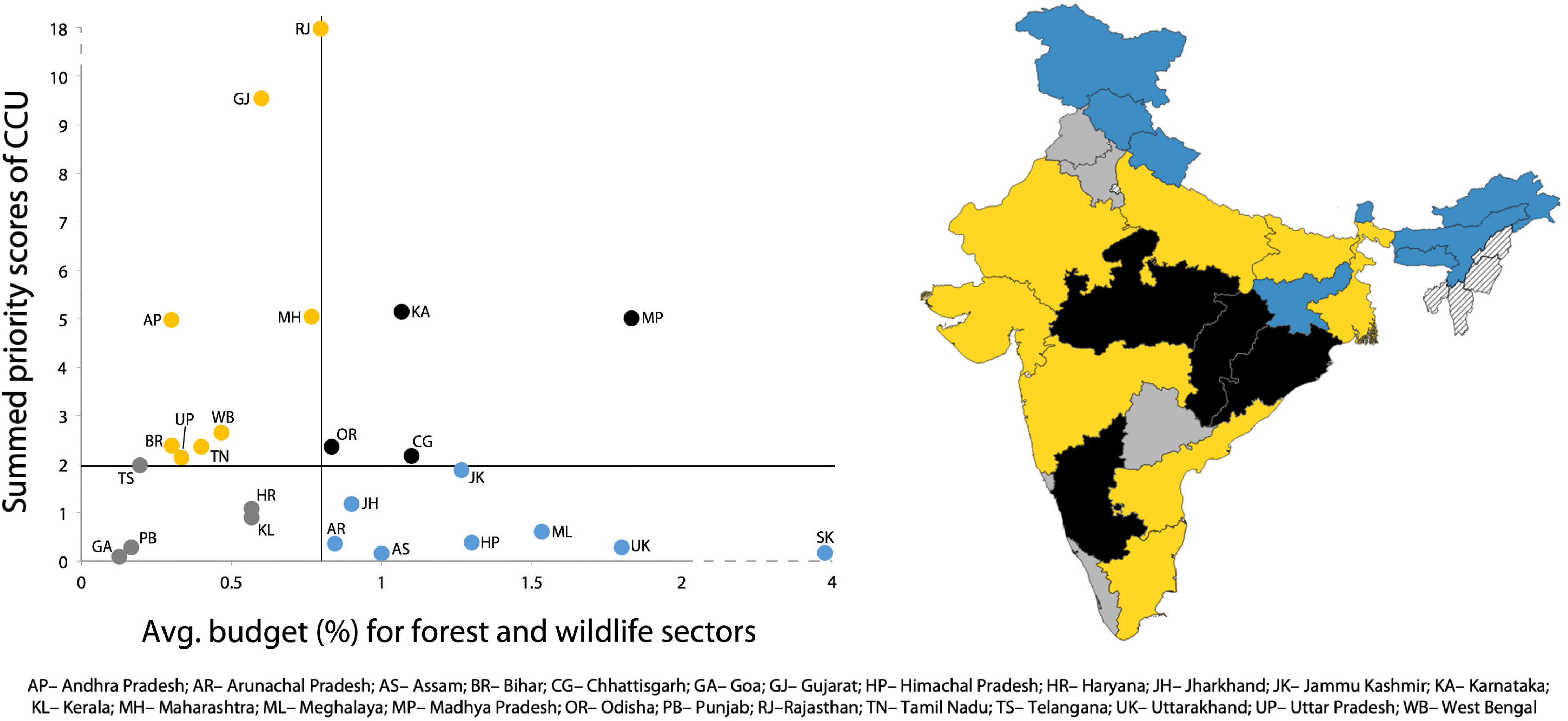
Summed CCU scores for each State in India, against the State’s average budgetary spending in wildlife and forest sectors (expressed as a proportion of annual State budget 2015– 2018). Vertical black lines on both axes are corresponding median values. Dots in black represent States with high CCU scores and higher budget allocation; yellow– high CCU scores, lower budget; grey– low CCU scores, low budget; blue– low CCU scores, higher budget. States with dashed lines do not have any sub-districts in the top 30% priority scores.

We extracted information on numbers and spatial locations of infrastructure projects across the country from multiple online databases of federal and State government ministries. A total of 1143 projects were implemented or commissioned in the last five years (roughly coinciding with the period considered for carnivore distribution assessment). We could correctly assign spatial locations to 1073 projects: 413 solar farms, 214 irrigation projects, 20 large mines, 205 large- and 221 small-hydropower projects. Of these, 41% overlapped with CCUs (Fig. 5), suggestive of high risks to carnivores and their habitats. Only 18% of the projects were in low conservation priority areas (sub-districts with bottom 30% priority scores), i.e., locations that ranked high in terms of projected human populations and poverty index.

**Figure 5.**
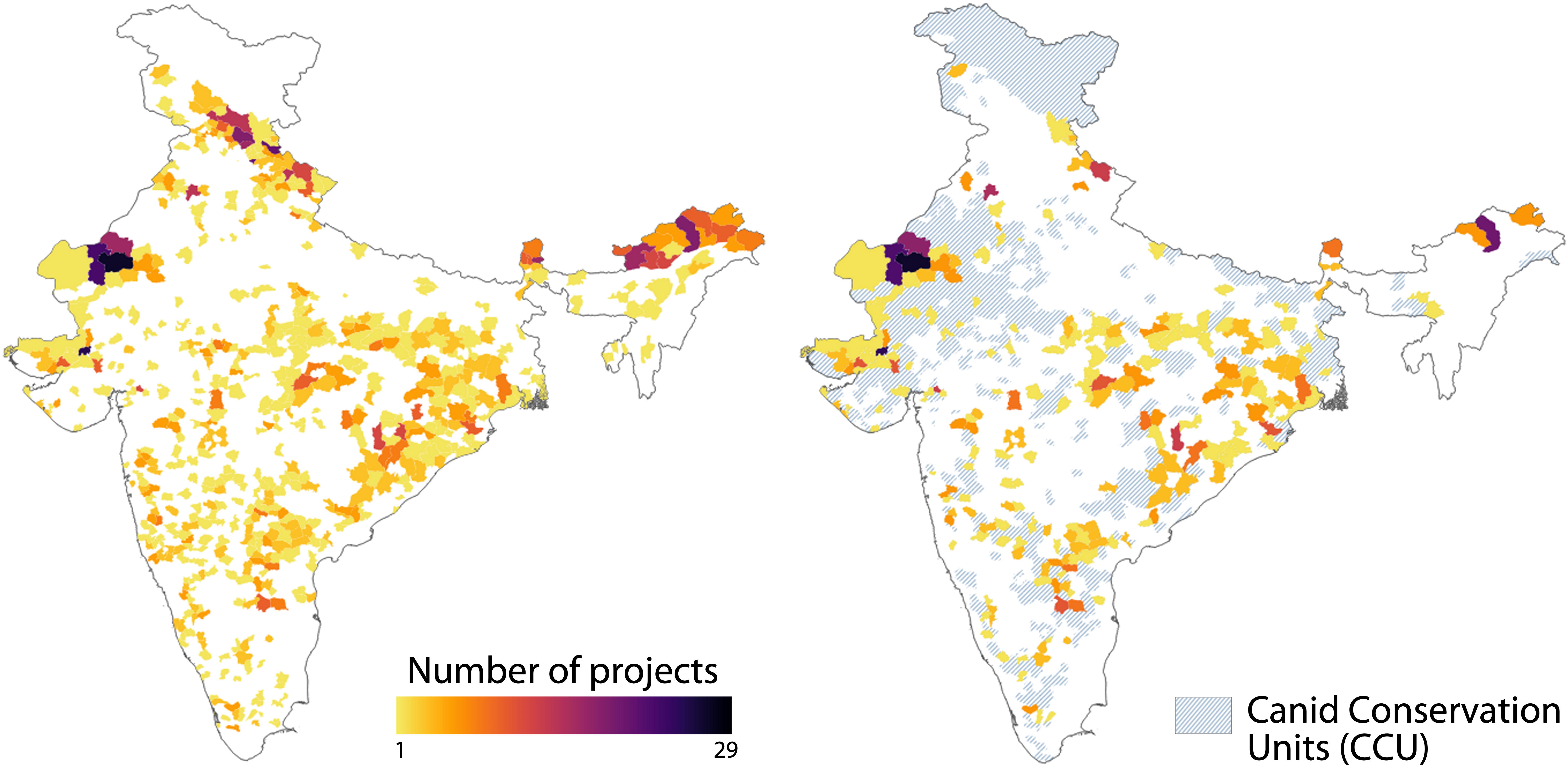
Infrastructure projects in key habitats. Left panel– Sub-district-wise numbers of infrastructure projects (irrigation projects, large mines, solar farms, large- and small-hydropower projects) implemented or commissioned in the past five years in India. Right panel– Sub-districts where infrastructure projects overlap with CCUs; dashed areas represent other CCUs with no projects or no available data on the key projects.

## Discussion

Given the oft-overlooked caveats with global-scale assessments, such as data inaccuracy or inadequacy and broad generalization of inferences (Ocampo-Peñuela et al. 2016; Norman & White 2019), our study demonstrates the utility of carefully designed large-scale investigations integrated with local knowledge and contexts. We present more nuanced distribution maps depicting species occurrences as probabilities within their Area of Habitat (Brooks et al. 2019; Fig. S2), rather than Minimum Convex Polygons that over- or under-estimate species ranges (Ocampo-Peñuela et al. 2016; Ramesh et al. 2017). Our approach also rectifies errors in current IUCN distribution maps; spatial mismatch between the two ranged from 167,897 sq. km for wolf (*Canis lupus pallipes* and *Canis lupus chanco*, combined) to 2,123,324 sq. km for red fox (*Vulpes vulpes* and *Vulpes vulpes pusilla*, combined). These findings are extremely relevant to past and future studies that use IUCN range maps for large-scale conservation assessments, reviews or syntheses. The focal species together represented multiple important habitats hitherto neglected in terms of conservation and management. Our estimates for Tibetan wolf and desert fox had lower reliability due to gaps in data (SI Appendix section 2), which we hope to address in future surveys.

About 5% of India’s 3.29 million sq. km land area is protected as National Parks and Wildlife Sanctuaries. Most of these are in forest habitats, owing to the country’s history of forest-centric protection regimes that stem from a British-colonial legacy of caching forestlands for timber extraction (see Ghosh-Harihar et al. 2019). Consequently, non-forest habitats that were either non-arable or unfit for timber production were termed “wastelands” (Singh et al. 2006). Decades of sidelining these habitats from mainstream conservation discourse has rendered them fragmented, isolated and degraded in most parts of the country (Vanak et al. 2015; Mondal et al. 2019). The systematic marginalization, compounded by other factors, resulted in the near-extinction of grassland-dependent species like the Great Indian Bustard *Ardeotis nigriceps*, Jerdon’s courser *Rhinoptilus bitorquatus*, and perhaps many others (Rahmani 2012). We foresee immense potential for expanding the conservation enterprise in India through demarcation of Conservation Reserves and Community Reserves, which currently cover *c*. 5,000 sq. km. Employing mechanisms such as creation of Village Reserves with equitable regulation can protect species and habitats without restricting sustainable use of resources in human-dominated landscapes. Our study also identifies locations where other logistically, economically, and socially feasible management options could be explored. If prioritized and carefully protected, the CCUs could potentially safeguard around 23,830 sq. km of grasslands, 99,950 sq. km of scrublands, 124,450 sq. km of open habitats or ravines, and 232,430 sq. km of forests– together with the current Protected Area network. This, of course, should not dissuade practitioners from prioritizing individual species or habitats in areas beyond CCUs for targeting conservation efforts.

Supporting the world’s second largest human population, India continues to witness rapid land-use changes, with land conversions being relatively easier for grasslands, scrublands and even agricultural lands (Baka 2014; Vanak et al. 2015; Kennedy et al. 2019). But ecological considerations rarely figure in current policies of landscape management and land-use planning. Instead, flawed notions of ‘compensating for damage’ through afforestation initiatives– when forests are cleared for infrastructure projects– are implemented in grasslands and scrublands (Vanak et al. 2015). Similar afforestation schemes globally promulgated for carbon sequestration and combating climate change (Bond et al. 2019) and exploiting these beleaguered habitats for industrial-scale solar farms, further increase their fragility while also bearing negative consequences for local communities (Yenneti et al. 2016; this study). The glaring knowledge gaps about these habitats impede our ability to comprehend their importance or plan their management (Ratnam et al. 2016). Our results add to the increasing evidence in literature that conservation of carnivores should necessarily include multi-use landscapes (including agricultural lands), considering their home range sizes and ecological requirements. Changes in crop type, transitions from seasonal to year-round cultivation, expansion of permanent irrigation, and intensification of pesticide use (impacting lower trophic levels) would create altered landscapes (UNCCD 2017), and potentially lead to local extinctions and/or shifts in carnivore communities (e.g., Majgaonkar et al. 2019). Future studies will need to undertake detailed investigations on these aspects to offer deeper ecological insights and augment our findings.

Carnivores have long held the fascination of wildlife managers, governments and the civil society (Treves & Karanth 2003; Smith et al. 2012). But they also present complicated conservation paradigms because of their negative interactions with people (Ripple et al. 2014; Montgomery et al. 2018); although, wild canids and hyenas in India are generally not associated with major threats to human life or property (Srivathsa et al. 2019). At present, wildlife monitoring in India predominantly involves national estimation of tigers and their prey, Asiatic lion, Asian elephant and one-horned rhinoceros populations. No protocol or framework exists for monitoring other species groups like wild canids, small carnivores, amphibians, riverine fauna, or ungulate herbivores in high-altitude or non-woodland habitats– a trend that appears to be commonplace for many species and habitats in developing countries (see Dornelas et al. 2014). Bridging some of these gaps, our work adds to a growing body of studies that leverage emerging opportunities in citizen science, social media and open source data for conservation monitoring (Newman et al. 2012; Ramesh et al. 2017; Toivonen et al. 2019).

Our focal species align themselves mid-way between obscurity and ubiquity, i.e., they are neither so rare that citizens are unaware of their existence, nor are they so abundant that calling for their conservation seems futile. And since these carnivores also occupy mid–high trophic levels, we believe they can serve as ideal flagships across important habitats (forests, alpine and arid/semi-arid grasslands, scrublands, ravines, open barren lands and deserts), which constitute ~41% of non-urban, non-agricultural areas in India. These results hold certain significance, since most recent studies determining global priority hotspots either exclude India altogether (Brum et al. 2017), or, include only forested areas in parts of Western Ghats and Eastern Himalayas (Brooks et al. 2006; Turner et al. 2007; Jenkins et al. 2013). Our aim is to build a monitoring system with real-time documentation of distribution and threat data on these carnivores, coupled with periodic analyses that would incrementally update the findings presented here. We anticipate the application of our approach to other poorly-studied taxa for which intensive resource allocation is not possible. This would complement government-driven monitoring programs with decentralized local efforts, fostering citizens’ involvement in scientific research while also garnering public support for conservation (Newman et al. 2012).

## Conclusion

The window of opportunity for implementing conservation initiatives in developing economies is becoming narrow. For spatial prioritization results to be relevant for local and federal governments, decision-support tools should incorporate human needs, politics and economic interests (Ban et al. 2013; Dickman et al. 2015). While our study does address these aspects, we recognize greater scope for refining and increasing the rigor of this exercise as we move forward. For instance, examining habitat connectivity and social thresholds of human–carnivore coexistence, quantifying region-specific threats and species-wise infrastructure impacts, tracking shifts in human populations towards urban centres and away from forest-dependent or agrarian livelihoods, and, smaller-scale assessments of costs and returns-on-investments, would all contribute towards adaptively building upon the results presented here. Nonetheless, we strongly argue for broadening current management notions of wilderness and natural habitats in India and recommend exploring and implementing alternative conservation approaches beyond forested Protected Areas. Certainly, executing these approaches would entail choosing habitat-specific flagships relevant to ecological, regional and social contexts, and developing monitoring techniques that can be co-opted by conservation managers and planners. In sum, our study represents an ‘ensemble approach’, combining a range of data sources, methods and analytical frameworks, and incorporates human population, poverty, infrastructure projects, and administrative potential and likelihood in conservation prioritization. We believe these are crucial for critically evaluating large-scale prioritization assessments, and for rethinking country-level conservation policies in India and other developing economies.

## Acknowledgements

We express gratitude to all the people who proactively participated and contributed data as part of our citizen-science surveys. We are also grateful to all the field experts in India who contributed to the threat assessment surveys. We thank Wildlife Conservation Society–India, IUCN Canid Specialist Group, Nature Conservation Foundation, Nature in Focus, Conservation India, IUCN Hyaena Specialist Group, R. Sreenivasan, V. Athreya and R. Chakravarty for their support. A.S was supported by the University of Florida, Wildlife Conservation Society’s Christensen Conservation Leaders Scholarship and Wildlife Conservation Network’s Sidney Byers Fellowship. G.A.P was supported by IDEA Wild. We thank M.A. Agnishikhe, A. Das, T. Kothawalla, D. Ganguly, K. Chauhan and A. Simon for their enthusiastic assistance in data compilation and processing. The authors received no funding for this study.

## Author contributions

A.S., I.M., S.S., P.S and G.A.P. designed the research; all authors performed the research and analyzed the data; A.S., I.M., P.S. and G.A.P. wrote the paper.

## Data availability

The data that support the findings of this study are available from the corresponding author upon reasonable request.

