## Supplementary material for "Opportunities for prioritizing and expanding conservation enterprise in India using carnivores as flagships": SI Appendix

**This file includes:**

Supplementary text
Figures S1 to S2
Tables S1 to S3
SI References

**Supplementary Text**

### SI Appendix: Section 1

### A. Phase 1: Citizen Science

Phase 1 of the project included records obtained through citizens' self-reported information. Sources include (a) survey form on a dedicated website, (b) information received through emails or phone calls and (c) information submitted through filled-in excel spreadsheets sent through emails and (d) information obtained from other citizen-science web portals.

**1. Survey form**

Single entry: A survey form was designed on the website (www.wildcanids.net) hosted on www.weebly.com. The form was designed to obtain information on single/one-off sightings by respondents. Data pertain to sighting records from 1 Jan 2015 to 31 Dec 2018. The format is given below:

**Respondent's particulars:**
Name:
Email:
Profession:

**1. Species**

- Dhole (*Cuon alpinus*)
- Jackal (*Canis aureus*)
- Indian Wolf (*Canis lupus pallipes*)
- Tibetan Wolf (*Canis lupus chanco*)
- Indian Fox (*Vulpes bengalensis*)
- Desert Fox (*Vulpes vulpes pusilla*)
- Red fox (*Vulpes vulpes*)
- Tibetan Fox (*Vulpes ferrilata*)
- Striped Hyena (*Hyaena hyaena*)

**2. Geographic location** (Location name, geographic coordinates)

**3. Habitat**

- Forest
- Scrub
- Rocky outcrop
- Grassland
- Peri-urban/ Rural
- Urban cities
- Agricultural field

Desert
High altitude plains
High altitude mountain
Coast and Mangroves

*Habitat description:*

**4. Type of sighting (Inside PA or Outside PA)** (this did not include reserve forests)
**5. Date** and **Time** [Day (7am to 5pm)| Night (7pm to 5am)| Dawn/Dusk (5am to 7pm and 5pm to 7pm)]
**6. Number of individuals**
**7. Number of young ones seen**
**8. Disease** Y/N
Description: (Eg. salivating profusely, mange/scabies/skin disorder...etc)
**9. Dead or Alive;** If dead, reason (this included roadkill, hunting/ poaching/ retaliatory killing/ Unknown/Other)
**10. Crop loss/ Livestock attack/ Human Attack**(with Description)
**11. Other interactions with humans** (e.g. provisioning food, kept as pets, feeding on garbage etc.)
**12. Other comments:** (e.g. was it an adult or a juvenile. Or adult with juveniles. In winter, many photos appear on social media of fox families, hyena families, etc.)

Multi entry: Forms were designed separately for those respondents with multiple records from one or more locations over a certain period of time. These were called “multi-entry response forms” and a Google survey form was designed for this purpose.
The information collected from each respondent was:
**Name:**
**Email:**
**Profession:**
**Species ID:**
**Time duration within which sightings occurred (range- MM/YY to MM/YY):**
**Type of sighting: (camera trap/direct sighting/other)**
**Geographic location:**
**Frequency of sighting (daily/monthly/infrequent):**
**Highest number of individuals seen:**
**Highest number of juveniles seen:**
**Habitat type:**
**Sighting location wrt to Protected Area:**
**Time of the day:**
**Death:**
**Diseases:**
**Crop loss; Livestock/Human injury:**

**Other interactions with humans:**
**Other Comments:**

In case of geographic location, we intended to collect information at the scale of a sub-district or a resolution lower than that (e.g., village name). We converted the location information to lat/long coordinates. These were then associated with sub-district codes in ArcGIS. We verified each record for the following:

1. Whether the species sighting was from a plausible location
2. Whether the location mentioned was consistent with other descriptions by the respondent: inside/outside PA, habitat type, etc.
3. The type of sighting: whether it was a direct sighting or not, since some people submitted records of vocalization, scat and pugmarks.
4. Requesting respondents to share photographs of sightings

If the sightings panned for more than a month, we quantified these based on the number of months. We deleted records if: the sightings were older than 2015, it was an indirect sign, the person did not respond to verification queries, or if it was a repeat record from the single entry form. We used the following standardization procedure to match the months with frequency of sightings:
(a) 1 month period and infrequent- once
(b) 2 month period and infrequent- once per month
(c) 2-4 months- assign a sighting for first and last month.
(d) 5 months and more- first and last month and one in between (for each year)

2. Eliciting survey responses

   a. An email blast was sent to a mailing list compiled by project team members. Individual mails were also sent to potential respondents who would submit multi-entry records.
   b. Social media posts were put up to announce the launch of the project. This was followed by daily/weekly posts to encourage citizens' participation.
   c. Social media promotions were aided by IUCN Canid SG page, Sanctuary Asia, Conservation India, etc..

3. Validation

   a. Data were downloaded at the end of each week in a comma-separated format (csv) in species-specific folders.
   b. The validation process involved checking for credibility of the record based on the information provided in the survey form (based on location, time of sighting, number of individuals, other comments).

c. This process also involved updating geographic coordinates of location, using the name of village, PA provided by the respondents (for records where the contributor had not included the same during submission).

*Follow-up:*

a. Follow-up emails were sent to contributors in case of any query. These included errors in dates, ambiguous or wrong locations, mismatch in location name and habitat type, mismatch in species ecology and habitat, or pack/group size, etc.
b. The updates on these were closely monitored and changes were made immediately after receiving a response.
c. In cases where the contributor did not respond back, some of these queries were internally clarified based on other details provided.
d. Follow up was also done for those records with interesting details (species-interactions, diseases, persecution/retaliatory killing, etc.), and requests for photographic records were made.

4. Mapping: Periodically, species-wise geographic coordinates were extracted and mapped. This facilitated spatial validation, as well as in making maps for the interim reports with survey participants.

### B. Phase 2: Web surveys

Phase 2 of the project involved systematic Internet searches from non-academic records of the focal species from social media websites, online photo-repositories and reliable blog articles or natural history photo-stories. Data pertained to sighting records from 1 Jan 2015 to 31 Dec 2018.

***Sources for Phase 2:***

**Facebook**

a. Keyword searches in the general search tab were done using single and plural versions of all species (e.g., "Indian wolf" and "Indian wolves"), and also the alternate names (e.g., "Desert fox" and "white footed fox"/ "white-footed fox"). A more targeted search was done in specific groups/pages using State-wise local/regional names of the species (e.g., "siyar" for jackal).

b. For each photograph located in this manner, the details of the location and the date of sightings, as well as other relevant details available were recorded. If the required information was not available in the image or image caption, a follow up with the photographer(s) either through commenting on the post itself or sending them a private message was done.

c. The second set of searches was more targeted, within select pages and groups. It followed the same procedure as above, but confined *within* specific pages/groups. The searches were done using the search tab within the group. We selected the "Sort by: Most recent" option to search posts in chronological order. Some groups that were targeted were: Sanctuary Asia, Saevus Wildlife India LLP, CLaW, Canidae and Hyaenidae of India, Nature in Focus, Mammals of Indian Subcontinent.

**Instagram**

Instagram searches followed the same protocol as that of Facebook. However hashtags were used along with the keywords (e.g. "#Indianwolf" and "#Indianwolves"). Considering many profiles on Instagram undertake paid promotions (photographers pay a page to feature their photographs), the original image was traced back to the photographers' own page to check for details. Follow up messages similar to that on Facebook were sent to obtain information that were not available in the image or image caption.

**India Nature Watch (INW)**

The India Nature Watch Portal is the oldest and perhaps the largest image-sharing website for wildlife of India. Tagged keywords were used to locate photographs of focal species. Since the database is vast, only those records where caption includes month/year and exact location of sighting were considered. Follow up with the photographer for details was avoided unless it was a rare species or an unusual location. Weblink: http://www.indianaturewatch.net

**Nature in Focus (NiF)**

Keyword searches were done to get information on the focal species using the image gallery "Hive" managed by Nature in Focus. Photos were usually associated with the month/year and location details. And in cases where these details were unavailable, the photographer was contacted. Weblink: https://hive.natureinfocus.in

**Mammals of India**

Mammals of India is a web portal (part of Biodiversity Atlas – India) hosted by National Centre for Biological Sciences (NCBS). Search for individual species was done in the drop down menu and information on location, and month/year of the record(s) was extracted for the focal species. Weblink: http://www.mammalsofindia.org

**Internet articles and blogs**
Image searches on google were carried out using the names of the species along with the word "India". The original article or blog was located by clicking on the species images. This search included websites such as Flickr, YouTube and Twitter (among others)

***Steps followed while entering Phase 2 data:***

1. Separate excel files were maintained for each species (9 files).
2. Month/year was entered in MM/YYYY format
3. Sub-district name was entered under "Location" and the State was mentioned in the "State" column. Google maps, internet searches or shapefiles were cross-verified to locate the sub-district name.
4. The geographic coordinates of the sub-district centroid was included in the 'Latitude' and 'Longitude' column.
5. Name of the person who has posted/uploaded the image or blog article was added under the 'Uploader name' column.
6. In the "Source" column, the source site was mentioned. [Options: "Facebook", "Instagram", "INW", "NiF", "NCBS", "Flickr", or "Internet article"].
7. Additional information from image/image caption (pack size, hunting/feeding info, threats etc.). **This was an optional column.**
8. The web link for the photograph or website where the image can be accessed was added to the "Source Link" column.

### C. Phase 3: Literature surveys

Phase 3 was carried out by the project team members, and involved tapping into scientific publications that featured/mentioned the details of the focal species.

a. A keyword search which included the common name and scientific name, followed by "India" was done using Google scholar and Web of Science.
b. Scientific articles published from 2015–2018 were considered, out of which only those data-points were used which were collected from January 2015 onwards.
c. Information was recorded in a specific data format. The citation of the article, and the journal name was entered in the "citation column ("Author (year): Journal Name").
d. This was followed by the duration of the study (in MM/YYYY format). In those cases where study started before 2015, the duration of the study was considered from January 2015, and the respective data-points were taken from the same time period. The actual/full duration of the study was mentioned in the Remarks column for reference.
e. The method of study used was mentioned. This included direct sightings, camera traps, sign surveys, interviews, secondary information. Interview-based studies

were not included to obtain data-points, but the area of study was used to identify plausible sub-districts where the species could be present.

f. This followed the entry of “Type of estimate” which included abundance index, occupancy, presence record, or any other. Type of estimate was carefully examined and based on the details in the article as well as credibility of the article/journal decisions were made whether to include data points from the article, or not.
g. The habitat type/s was recorded based on the study area description in the article.
h. The sighting location (PA/non-PA/Both) was also recorded based on the study area description in the paper.

Data processing: Data points were further broken down for each month and sub-district based on the information available.

a. If the species of interest was continuously monitored/encountered during the study, the data point with presence information was attributed to all the months within the study period for respective sub-districts.
b. In studies where the species was opportunistically encountered, only the months in which it was seen/photo-captured/recorded was considered as a data-point for the sub-district.
c. In cases where the exact months could not be determined directly, ecological clues (e.g., breeding, denning, etc.) were used, or, the data were discarded.

### SI Appendix: Section 2

**Description of the occupancy modeling process:** For each species, we identified a set of land-use land-cover (LULC), ecological/climatic and anthropogenic variables, predicted to influence their occupancy patterns. The full list of these variables, categories, descriptions and sources are provided in Table 2. All variables were processed at the sub-district level and z-transformed prior to analyses. We also checked for cross-correlations between variables and refrained from using highly correlated variables (Pearson's correlation $r>|0.6|$) in the same model.

In the first step, we retained an intercept-only formulation for *psi* and compared the relative fit of singular and additive model combinations for *p*. We modeled detectability at the site-level (for all species) as a function of four factors: average distance to four closest big cities, density of roads/railways in the sub-district (data sourced from Open Street Map OSM hosted on Geofabrik; http://download.geofabrik.de), presence/absence of Protected Area, and the size of the sub-district. We expected these factors to influence people's access to sub-districts, together with our ability to detect records of species presence. In the second step, we retained the variable(s) for *p* from the top-ranked model in step 1, and compared the relative fit of models with singular effects for *psi*. Model comparisons were based on Akaike's Information Criterion, corrected for small samples sizes (AICc; Burnham & Anderson 2002). To maintain parsimony, we used only up to four additive models for each species: one with a combination of LULC variables, one with ecological/climatic variables, one with anthropogenic variables and one with four top-ranked variables from the model runs with singular variables.

We checked for goodness-of-fit for each species using parametric bootstrapping, as described in MacKenzie & Bailey (2004). The $\chi^2$ statistic for the top-ranked model for all species, except Tibetan wolf and desert fox, was lower than the average of the simulated test statistic ($p < \hat{c}$), indicating no over-dispersion in the data. The final estimates for *psi* for each species were derived from averaging across all models whose cumulative AICc weight added to >0.90. For each species, we calculated the "area of occupancy" by multiplying its estimated occupancy probability in every sub-district with the extent of species-specific habitats (in sq. km) in the sub-district. These estimates were summed across all sub-districts within each species' plausible range to arrive at the corresponding area of occupancy in the country.

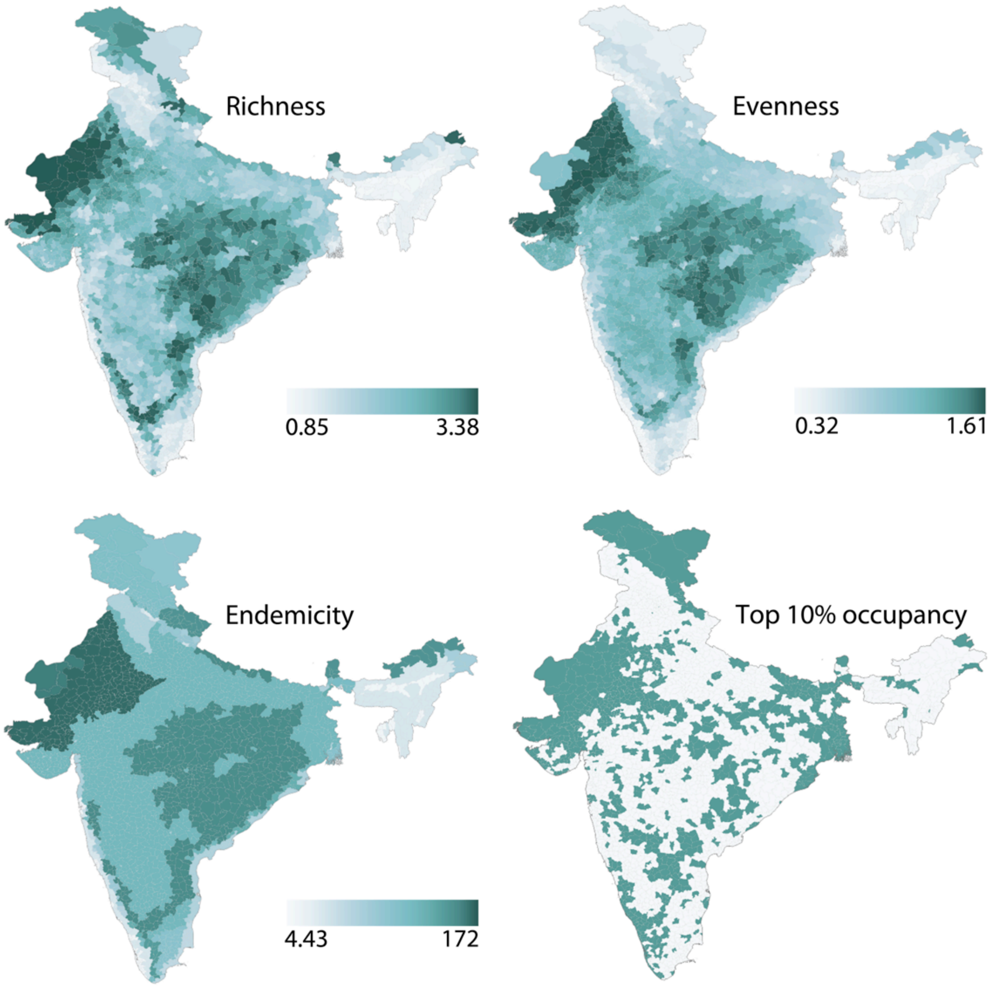

**Figure S1.** Sub-district level species richness (weighted by species' conservation score), evenness (calculated as Shannon-Weiner index using species' area of occurrence), combined endemicity scores, and sub-districts with top 10% occupancy probability (darker shade) for at least one of the focal species.

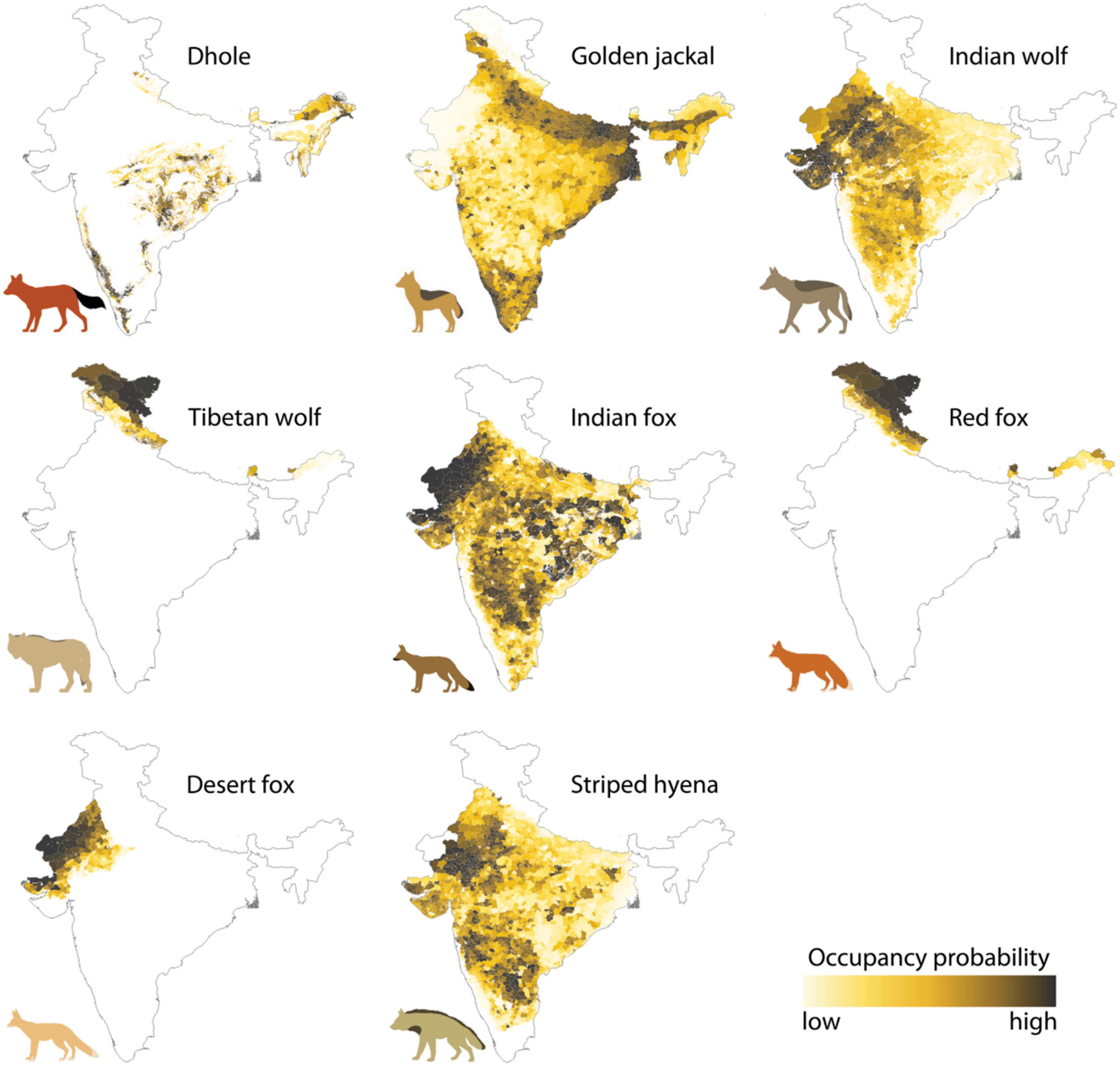

**Figure S2.** Spatial occupancy probabilities for focal carnivore species within the extent of species-specific plausible habitats.

**Table S1.** Descriptions of categories and data sources for explanatory variables used for modeling carnivore distributions in India

| Sl. | Category | Description | Source; Year |
|---|---|---|---|
| 1 | Land-use land cover (LULC) | 153 LULC categories were combined into 12 classes: temperate forest, tropical dry forest, tropical dry forest, scrubland, grassland, open (barren) areas, ravines, high altitude plains, high altitude mountains, coastal/mangrove, agriculture, and production agroforests | Indian Institute Remote Sensing, Govt. of India; http://bis.iirs.gov.in; 2015 |
| 2 | Habitat extent | For each species, we selected subsets of relevant LULC types and calculated the total area (sq.km) in every sub-district within their plausible range. | Indian Institute Remote Sensing, Govt. of India; http://bis.iirs.gov.in; 2015 |
| 3 | Human settlements | Extent of human settlements was taken from the LULC dataset described above | Indian Institute Remote Sensing, Govt. of India; http://bis.iirs.gov.in; 2015 |
| 4 | Rocky barren areas | Area covered by rocky outcrops and escarpments were calculated from Wasteland map of India | Resourcesat-1 LISS III; http://bhuvan5.nrsc.gov.in/bhuvan/wms; 2006 |
| 5 | Precipitation | Monthly rainfall data was downloaded for September 2015–August 2016. Average monthly rainfall for each sub-district was summed across 12 months | Tropical Rainfall Measuring Mission TRMM imagery NASA Goddard Earth Sciences DISC, https://earthobservatory.nasa.gov/global-maps/; 2015–2016 |
| 6 | Elevation | Void-filled Digital Elevation Model data was downloaded and average elevation calculated for each sub-district | USGS: HydroSHEDS; https://hydrosheds.cr.usgs.gov/index.php; 2005 |
| 7 | Ruggedness | Ruggedness index (terrain heterogeneity) was calculated for the elevation raster using 'terrain analysis' tool on QGIS, and averaged for each sub-district | USGS: HydroSHEDS; https://hydrosheds.cr.usgs.gov/index.php; 2015 |
| 8 | Wild prey | Wild prey index was calculated for dhole and Indian wolf based on published literature. Each sub-district was assigned a value between 0 (absence) to 4 (high [illegible] | Karanth et al. (2009) |

| | | | |
|---|---|---|---|
| 9 | Livestock population | Livestock (cows, buffalo, goats, sheep) population data were compiled from government census records | All India Livestock Census 2012, Govt. of India; http://dahd.nic.in; 2012 |
| 10 | Dog population | Number of domestic and free-ranging dogs were compiled government census records | All India Livestock Census 2012, Govt. of India; http://dahd.nic.in; 2012 |
| 11 | Human population | Estimated human population for the year 2015 (using baseline population from 2011 census by Govt. of India). Population density calculated for each sub-district | The Gridded Population of the World, Version 4 (GPWv4). NASA Socioeconomic Data and Applications Center (SEDAC); https://sedac.ciesin.columbia.edu; 2015 |
| 12 | Linear infrastructure | Linear infrastructure (roadways and railways) data were downloaded and processed at the sub-district level. Final value for each sub-district was the total length (in km) divided by area of sub-district | OpenStreetMap (OSM) Project; http://download.geofabrik.de; 2018 |
| 13 | Protected Areas | Data on Protected Areas (National Parks and Wildlife Sanctuaries) of India were downloaded in vector format from WDPA and updated using from WII. For each sub-district we calculated area and proportion covered by Protected Area boundaries | World Database on Protected Areas (WDPA), ENVIS Centre on Wildlife & Protected Areas (Wildlife Institute of India, WII); https://www.protectedplanet.net and http://wiienvis.nic.in/Home.aspx; 2018 |

**Table S2.** Descriptions of categories, relative weights and data sources for input variables used for spatial prioritization analysis. Carnivore diversity indices were ranked highest, followed by key habitats. Predicted human population and poverty index were treated as 'costs' and therefore assigned negative weights

| Sl. | Category | Description | Relative Weight | Source; Year |
|---|---|---|---|---|
| 1 | Top 10 percent | Sub-districts with top 10% occupancy probabilities for one or more species. Selected sub-districts were assigned code '1' and rest as '0'. | 9 | Derived from occupancy analysis |
| 2 | Endemicity | Endemicity for each species was calculated as the inverse of the area it occupied across the country. Endemicity values were summed across all species occurring in each sub-district | 8 | Derived from occupancy analysis |
| 3 | Evenness | Shannon-Weiner index calculated using species' area of occurrence in lieu of number of encounters. The final index is the summed values for all species occurring in each sub-district | 7 | Derived from occupancy analysis |
| 4 | Richness | Richness was calculated for each sub-district as the weighted sum of individual species' occupancy probabilities. Individual probabilities were weighted by conservation scores (more threatened species received higher weights) | 6 | Derived from occupancy analysis |
| 5 | Habitat area | Key habitats were weighted based on degree of vulnerability to change/conversion/diversion. Forest=1, agriculture=2, open habitats and ravines=3, scrublands=4 and grasslands=5 | 1–5 | Indian Institute of Remote Sensing, Govt. of India; http://bis.iirs.gov.in; 2015 |
| 6 | Human population | Predicted human population for 2020; processed for each sub-district | -1 | NASA Socioeconomic Data and Applications Center (SEDAC); https://sedac.ciesin.columbia.edu; 2018 |
| 7 | Poverty index | Poverty head count (%) data from were extracted from government census records and re-processed at the sub-district level | -1 | Mohanty et al. (2016) |

**Table S3.** Model comparisons for estimating probability of presence (*psi*) and detectability (*p*) of the focal carnivore species. For each species, the first set of models are for estimating detectability while the covariate structure for occupancy parameter is held constant (intercept-only formulation); the second set of models are for estimating occupancy probability, while retaining the top-ranked variable(s) for detectability. Up to top five models for *p* and up to top ten models for *psi* (both based on AICc ranks) are presented for each species. Abbreviation codes are in the footnote.

| **species: dhole** | **params.** | **AICc** | **ΔAICc** | **AICc weight** | **cuml. weight** |
|---|---|---|---|---|---|
| **Model comparisons for *p*** | | | | | |
| *psi, p(rpres+area+city+infa)* | 6 | 740.93 | 0 | 0.32 | 0.32 |
| *psi, p(rpres+area+city)* | 5 | 741.4 | 0.47 | 0.25 | 0.57 |
| *psi, p(rpres+area+infa)* | 5 | 741.63 | 0.7 | 0.22 | 0.79 |
| *psi, p(rpres+area)* | 4 | 743.24 | 2.31 | 0.1 | 0.89 |
| *psi, p(rpres)* | 3 | 745.03 | 4.1 | 0.04 | 0.93 |
| **Model comparisons for *psi*** | | | | | |
| *psi(prey+resv+hpop+catl), p(covs)* | 10 | 712.57 | 0 | 0.99 | 0.99 |
| *psi(prey), p(covs)* | 7 | 722.27 | 9.7 | 0.01 | 1 |
| *psi(prey+pptn+rugg), p(covs)* | 9 | 725.92 | 13.34 | 0 | 1 |
| *psi(hpop+resv+catl+dogs), p(covs)* | 10 | 728.46 | 15.89 | 0 | 1 |
| *psi(resv), p(covs)* | 7 | 728.79 | 16.22 | 0 | 1 |
| *psi(hpop), p(covs)* | 7 | 738.92 | 26.35 | 0 | 1 |
| *psi(catl), p(covs)* | 7 | 739.56 | 26.98 | 0 | 1 |
| *psi(habt), p(covs)* | 7 | 740.77 | 28.19 | 0 | 1 |
| *psi(fcov), p(covs)* | 7 | 740.86 | 28.29 | 0 | 1 |
| *psi(rugg), p(covs)* | 7 | 740.86 | 28.29 | 0 | 1 |

| species: golden jackal | params. | AICc | ΔAICc | AICc weight | cuml. weight |
|---|---|---|---|---|---|
| **Model comparisons for *p*** | | | | | |
| *psi ,p(ifra+city+rpres+area)* | 6 | 2022.32 | 0 | 0.53 | 0.53 |
| *psi, p(ifra+city+area)* | 5 | 2022.89 | 0.57 | 0.40 | 0.94 |
| *psi, p(ifra+city+rpres)* | 5 | 2027.61 | 5.3 | 0.04 | 0.97 |
| *psi, p(ifra+city)* | 4 | 2029.92 | 7.61 | 0.01 | 0.98 |
| *psi, p (ifra+area)* | 4 | 2030.39 | 8.08 | 0.01 | 0.99 |
| **Model comparisons for *psi*** | | | | | |
| *psi(hpop), p(covs)* | 7 | 1999.86 | 0 | 0.39 | 0.39 |
| *psi(hpop+prod+ifra+pptn), p(covs)* | 10 | 1999.88 | 0.01 | 0.38 | 0.77 |
| *psi(prod+gsor), p(covs)* | 8 | 2002.12 | 2.25 | 0.13 | 0.9 |
| *psi(hpop+ifra+dogs), p(covs)* | 9 | 2002.76 | 2.90 | 0.09 | 0.99 |
| *psi(prod), p(covs)* | 7 | 2007.65 | 7.78 | 0.01 | 1 |
| *psi(pptn+rugg), p(covs)* | 8 | 2008.97 | 9.11 | 0 | 1 |
| *psi(pptn), p(covs)* | 7 | 2015.89 | 16.02 | 0 | 1 |
| *psi(ifra), p(covs)* | 7 | 2016.13 | 16.27 | 0 | 1 |
| *psi(sett), p(covs)* | 7 | 2016.39 | 16.53 | 0 | 1 |
| *psi(gsor), p(covs)* | 7 | 2017.36 | 17.50 | 0 | 1 |

| species: Indian wolf | params. | AICc | ΔAICc | AICc weight | cuml. weight |
|---|---|---|---|---|---|
| **Model comparisons for *p*** | | | | | |
| *psi, p(city)* | 3 | 722.63 | 0 | 0.32 | 0.32 |
| *psi, p(city+area)* | 4 | 723.83 | 1.2 | 0.17 | 0.49 |
| *psi, p(city+ifra)* | 4 | 724.37 | 1.75 | 0.13 | 0.62 |
| *psi, p(city+rpres)* | 4 | 724.63 | 2 | 0.12 | 0.74 |
| *psi, p(city+ifra+area)* | 5 | 725.47 | 2.84 | 0.08 | 0.82 |
| **Model comparisons for *psi*** | | | | | |
| *psi(prey+rugg+hpop+dogs), p(covs)* | 7 | 711.45 | 0 | 0.70 | 0.70 |
| *psi(prey+rugg+shot), p(covs)* | 6 | 714.37 | 2.92 | 0.16 | 0.86 |
| *psi(prey), p(covs)* | 4 | 716.22 | 4.77 | 0.06 | 0.92 |
| *psi(rugg), p(covs)* | 4 | 716.96 | 5.5 | 0.04 | 0.97 |
| *psi(dogs), p(covs)* | 4 | 720.19 | 8.73 | 0.01 | 0.97 |
| *psi(hpop), p(covs)* | 4 | 720.46 | 9.01 | 0.01 | 0.98 |
| *psi(hpop+ifra+dogs), p(covs)* | 6 | 722.10 | 10.65 | 0.003 | 0.99 |
| *psi(ifra), p(covs)* | 4 | 722.62 | 11.17 | 0.003 | 0.99 |
| *psi(sett), p(covs)* | 4 | 722.67 | 11.22 | 0.003 | 0.99 |
| *psi(habt), p(covs)* | 4 | 722.73 | 11.28 | 0.002 | 0.99 |

| species: Tibetan wolf | params. | AICc | ΔAICc | AICc weight | cuml. weight |
|---|---|---|---|---|---|
| **Model comparisons for *p*** | | | | | |
| *psi, p(area+city)* | 4 | 43.32 | 0 | 0.34 | 0.34 |
| *psi, p(area+ifra)* | 4 | 43.71 | 0.39 | 0.28 | 0.62 |
| *psi, p(area)* | 3 | 43.73 | 0.42 | 0.27 | 0.89 |
| *psi, p(city+area+ifra)* | 5 | 45.71 | 2.39 | 0.10 | 0.99 |
| *psi, p(city)* | 3 | 51.94 | 8.62 | 0.005 | 1 |
| **Model comparisons for *psi*** | | | | | |
| *psi(pptn), p(covs)* | 5 | 40.56 | 0 | 0.36 | 0.36 |
| *psi(ifra), p(covs)* | 5 | 43.01 | 2.45 | 0.10 | 0.46 |
| *psi(dogs), p(covs)* | 5 | 43.51 | 2.95 | 0.08 | 0.54 |
| *psi(habt), p(covs)* | 5 | 44.32 | 3.76 | 0.05 | 0.6 |
| *psi(elev), p(covs)* | 5 | 44.48 | 3.92 | 0.05 | 0.65 |
| *psi(pptn+ifra+dogs+habt), p(covs)* | 8 | 44.49 | 3.93 | 0.05 | 0.7 |
| *psi(sett), p(covs)* | 5 | 44.77 | 4.21 | 0.04 | 0.74 |
| *psi(shot), p(covs)* | 5 | 44.90 | 4.33 | 0.04 | 0.78 |
| *psi(rugg), p(covs)* | 5 | 44.90 | 4.34 | 0.04 | 0.82 |
| *psi(gsor), p(covs)* | 5 | 45.18 | 4.62 | 0.04 | 0.86 |

| species: Indian fox | params. | AICc | ΔAICc | AICc weight | cuml. weight |
|---|---|---|---|---|---|
| **Model comparisons for *p*** | | | | | |
| *psi, p(area)* | 3 | 903.02 | 0 | 0.28 | 0.28 |
| *psi, p(area+city)* | 4 | 904.03 | 1.01 | 0.17 | 0.45 |
| *psi, p(area+ifra)* | 4 | 904.05 | 1.03 | 0.17 | 0.61 |
| *psi, p(area+rpres)* | 4 | 904.29 | 1.27 | 0.15 | 0.76 |
| *psi, p(area+city+rpres)* | 5 | 904.94 | 1.92 | 0.11 | 0.87 |
| **Model comparisons for *psi*** | | | | | |
| *psi(habt+gsor+hpop+pptn), p(covs)* | 7 | 896.1 | 0 | 0.44 | 0.44 |
| *psi(habt), p(covs)* | 4 | 896.51 | 0.41 | 0.36 | 0.8 |
| *psi(gsor), p(covs)* | 4 | 900.45 | 4.35 | 0.05 | 0.85 |
| *psi(hpop), p(covs)* | 4 | 900.49 | 4.39 | 0.05 | 0.9 |
| *psi(hpop+dogs), p(covs)* | 5 | 902.3 | 6.2 | 0.02 | 0.92 |
| *psi(pptn), p(covs)* | 4 | 902.41 | 6.31 | 0.02 | 0.94 |
| *psi(dogs), p(covs)* | 4 | 903.20 | 7.1 | 0.01 | 0.95 |
| *psi(rugg), p(covs)* | 4 | 903.48 | 7.38 | 0.01 | 0.96 |
| *psi(sett), p(covs)* | 4 | 903.52 | 7.42 | 0.01 | 0.97 |
| *psi(pptn+rock+rugg), p(covs)* | 5 | 903.78 | 7.68 | 0.01 | 0.98 |

| species: red fox | params. | AICc | ΔAICc | AICc weight | cuml. weight |
|---|---|---|---|---|---|
| **Model comparisons for *p*** | | | | | |
| *psi, p(city)* | 3 | 142.82 | 0 | 0.26 | 0.26 |
| *psi, p(city+ifra)* | 4 | 143.54 | 0.72 | 0.18 | 0.43 |
| *psi, p(city+area)* | 4 | 144.03 | 1.21 | 0.14 | 0.57 |
| *psi, p(city+ifra+area)* | 5 | 144.46 | 1.64 | 0.11 | 0.69 |
| *psi, p(city+rpres)* | 4 | 144.63 | 1.81 | 0.10 | 0.79 |
| **Model comparisons for *psi*** | | | | | |
| *psi(elev+pptn), p(covs)* | 5 | 133.95 | 0 | 0.35 | 0.35 |
| *psi(elev), p(covs)* | 4 | 134.3 | 0.35 | 0.29 | 0.64 |
| *psi(pptn), p(covs)* | 4 | 134.77 | 0.82 | 0.23 | 0.87 |
| *psi(pptn+elev+dogs+gsor), p(covs)* | 7 | 137.18 | 3.23 | 0.07 | 0.94 |
| *psi(rugg), p(covs)* | 4 | 140.33 | 6.38 | 0.01 | 0.95 |
| *psi(gsor+hamt), p(covs)* | 5 | 140.82 | 6.87 | 0.01 | 0.96 |
| *psi(gsor), p(covs)* | 4 | 141.49 | 7.54 | 0.01 | 0.97 |
| *psi(hamt), p(covs)* | 4 | 141.83 | 7.88 | 0.01 | 0.98 |
| *psi(agri), p(covs)* | 4 | 142.11 | 8.16 | 0.01 | 0.98 |
| *psi(dogs), p(covs)* | 4 | 142.19 | 8.24 | 0.01 | 0.99 |

| species: desert fox | params. | AICc | ΔAICc | AICc weight | cuml. weight |
|---|---|---|---|---|---|
| **Model comparisons for *p*** | | | | | |
| *psi, p(area)* | 3 | 87.64 | 0 | 0.2 | 0.2 |
| *psi, p(area+ifra)* | 4 | 87.68 | 0.04 | 0.19 | 0.39 |
| *psi, p(area+ifra+rpres)* | 5 | 87.73 | 0.09 | 0.19 | 0.58 |
| *psi, p(area+rpres)* | 4 | 87.87 | 0.23 | 0.18 | 0.76 |
| *psi, p(area+ifra+city+rpres)* | 6 | 89.34 | 1.7 | 0.09 | 0.85 |
| **Model comparisons for *psi*** | | | | | |
| *psi(habt+gsor+ohab+ifra), p(covs)* | 7 | 79.87 | 0 | 0.57 | 0.57 |
| *psi(habt), p(covs)* | 4 | 80.69 | 0.82 | 0.38 | 0.95 |
| *psi(gsor+ohab), p(covs)* | 5 | 86.21 | 6.34 | 0.02 | 0.97 |
| *psi(gsor), p(covs)* | 4 | 88.23 | 8.37 | 0.01 | 0.98 |
| *psi(ohab), p(covs)* | 4 | 88.49 | 8.62 | 0.01 | 0.99 |
| *psi(ifra), p(covs)* | 4 | 89.64 | 9.77 | 0.004 | 0.99 |
| *psi(agri), p(covs)* | 4 | 89.64 | 9.77 | 0.004 | 1 |
| *psi(hpop), p(covs)* | 4 | 89.64 | 9.77 | 0.004 | 1 |

| species: striped hyena | params. | AICc | ΔAICc | AICc weight | cuml. weight |
|---|---|---|---|---|---|
| **Model comparisons for *p*** | | | | | |
| *psi, p(area)* | 3 | 871.26 | 0 | 0.32 | 0.32 |
| *psi, p(area+ifra)* | 4 | 872.56 | 1.3 | 0.16 | 0.48 |
| *psi, p(area+ifra+city)* | 5 | 873.06 | 1.8 | 0.13 | 0.61 |
| *psi, p(area+city)* | 4 | 873.11 | 1.86 | 0.12 | 0.73 |
| *psi, p(area+rpres)* | 4 | 873.22 | 1.96 | 0.12 | 0.85 |
| **Model comparisons for *psi*** | | | | | |
| *psi(rock), p(covs)* | 4 | 868.03 | 0 | 0.29 | 0.29 |
| *psi(prod), p(covs)* | 4 | 869.83 | 1.8 | 0.12 | 0.40 |
| *psi(gsor), p(covs)* | 4 | 870.25 | 2.22 | 0.09 | 0.50 |
| *psi(rock+prod+gsor+dogs), p(covs)* | 7 | 870.35 | 2.32 | 0.09 | 0.59 |
| *psi(prod+gsor), p(covs)* | 5 | 870.43 | 2.40 | 0.09 | 0.68 |
| *psi(dogs), p(covs)* | 4 | 871.04 | 3.01 | 0.06 | 0.74 |
| *psi(hpop), p(covs)* | 4 | 871.75 | 3.72 | 0.04 | 0.78 |
| *psi(dfor), p(covs)* | 4 | 872.25 | 4.22 | 0.03 | 0.82 |
| *psi(dogs+hpop), p(covs)* | 5 | 872.26 | 4.23 | 0.03 | 0.85 |
| *psi(sett), p(covs)* | 4 | 872.59 | 4.56 | 0.03 | 0.88 |

Abbreviations: fcov– combined forest cover; dfor– tropical dry forests; tfor– temperate forests; gsor– grasslands, scrublands, open (barren) habitats and ravines; ohab– open (barren) habitats; agri– agricultural areas; prod– production agroforests; hamt– high altitude mountains; habt– combined area of plausible habitats (species-specific); rock– rocky outcrops and escarpments; pptn– annual precipitation; rugg– terrain ruggedness; prey– wild prey index; elev– elevation; catl– cattle population; shot– sheep and goat population; lstk– livestock (cattle, sheep and goat) population; hpop– human population; dogs– free-ranging dog population; sett– human settlements; ifra– density of roads and railways; resv– extent of Protected Areas; area– area of sub-district; city– average distance to closest four big cities; rpres– presence/absence of Protected Area in the sub-district; covs– top-ranked variable(s) from model comparisons for *p*
